# Single-molecule sequencing maps replication dynamics across the fission yeast genome, including centromeres

**DOI:** 10.1101/2025.07.16.665067

**Authors:** Isabel Díez-Santos, Sathish Thiyagarajan, Anna M. Rogers, Adam T. Watson, Antony M. Carr, Conrad A. Nieduszynski

## Abstract

DNA replication dynamics in fission yeast remain incompletely understood, particularly in repetitive regions such as the centromeres and the rDNA. Here, we establish DNAscent—a nanopore sequencing method that detects BrdU-labelled nascent DNA and infers replication dynamics—to map replication at single-molecule resolution in fission yeast. For the first time, we have identified thousands of replication forks, as well as initiation, termination and pause events, on single sequenced molecules across the whole genome. This high coverage allowed us to identify replication patterns in poorly characterised regions. In the rDNA, we detect strand-specific pausing at replication fork barriers. At the mating-type locus, we find the most frequent pause site outside the rDNA. At centromeres, we find that replication initiation predominantly occurs in the outer repeats, while termination localises to central regions and that, only in centromere 2, there is an enrichment in pauses at the centromeric tRNAs. This work establishes a powerful single-molecule method for studying replication dynamics in fission yeast and provides insights into replication across repetitive regions that constitute a significant portion of the genomes of more complex organisms.

## INTRODUCTION

DNA replication is a fundamental process in all organisms by which the genome is accurately duplicated. In eukaryotes, DNA replication initiates at multiple sites—termed origins—across each chromosome. From these sites, bidirectional replication forks proceed until they terminate when meeting opposing forks. During this process, replication may pause due to obstacles such as protein barriers or DNA lesions. Perturbed forks or insufficient replication initiation events can be a source of genome instability (1–3). Therefore, significant effort has gone into developing methods to study the sites of replication initiation and the kinetics of replication fork progression.

Much of our understanding of replication dynamics—particularly in yeasts such as budding yeast (*Saccharomyces cerevisiae*; *S. cerevisiae*) and fission yeast (*Schizosaccharomyces pombe*; *S. pombe*)—has come from population-based, short-read genomic studies (4). In *S. pombe*, early genomic studies identified replication origin locations via chromatin immunoprecipitation (ChIP) of origin-associated proteins or the increase in DNA copy number at active origin sites (5, 6). More recently, a method termed polymerase usage sequencing (Pu-seq) identified not only replication initiation, but also termination and fork direction with high resolution (7). Pu-seq locates ribonucleotides erroneously incorporated into DNA by mutant replicative polymerases. Since polymerase ε and polymerase δ primarily synthesise the leading and lagging strands, respectively, the relative usage of these polymerases can be used to infer fork direction, and by extension, sites of initiation and termination. These methods rely on short-read sequencing and therefore cannot resolve the events occurring in complex repetitive areas such as the rDNA and centromeres. Moreover, they offer a population-level view of replication dynamics and fail to capture cell-to-cell variation.

Single-molecule and single-cell methods address the issue of cellular heterogeneity, but single-cell approaches often lack the resolution to precisely define replication events (8, 9). In fission yeast, DNA combing is the main single-molecule method that has been used to visualise major aspects of replication dynamics (10, 11). Advances in DNA combing allowed the analysis of ultra-long DNA molecules (>1 Mb) and provided information on replication dynamics in complex, repetitive areas such as the rDNA, centromeres and telomeres.

However, DNA combing relies on antibody-based detection of nucleotide analogues, which limits its spatial and temporal resolution and makes it challenging to map replication events precisely. Recently, an alternative single-molecule method, DNAscent, was developed to provide genome-wide, long single-molecule information on replication fork direction, initiation and termination sites, and fork pausing events (12–14). This technique was established in budding yeast (12), and has been successfully applied in cultured human cells (15, 16), and protozoan parasites (17, 18). DNAscent detects synthetic nucleotide analogues, such as bromodeoxyuridine (BrdU), incorporated into nascent DNA strands during replication (Figure 1A). Varying BrdU incorporation levels with respect to time allowed identification of where replication initiates and terminates, fork direction and sites of fork pausing.

**Figure 1:**
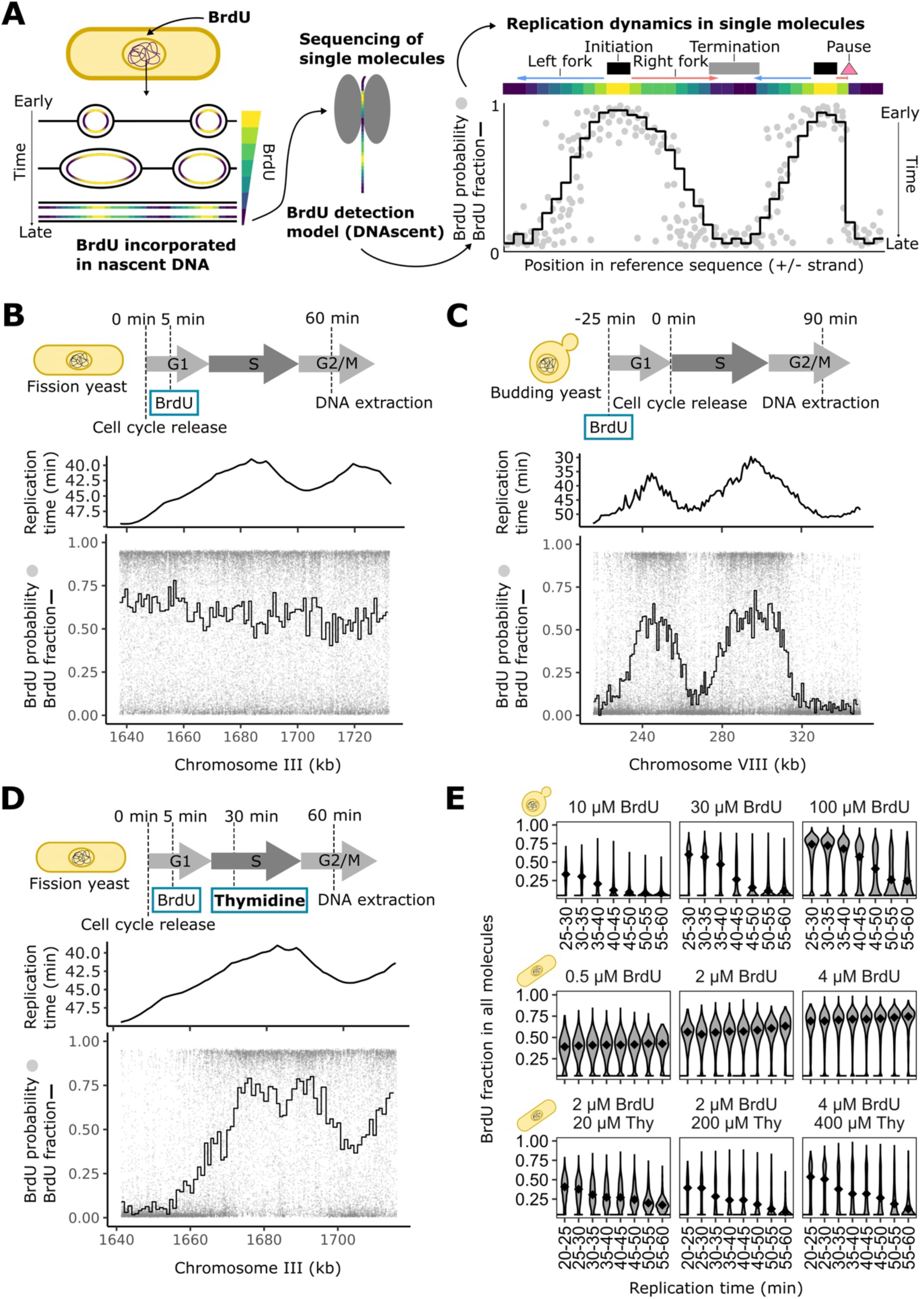
Levels of BrdU incorporation in fission and budding yeast. A) Overview of the methodology for detection of BrdU and inference of DNA replication dynamics on single molecules. Cells are grown in the presence of the nucleoside analogue BrdU, which is incorporated into DNA in place of canonical thymidine. Then, DNA is extracted and nanopore sequenced. The resulting reads are analysed with a model that detects BrdU (DNAscent; see Methods). The density of BrdU along the sequenced DNA molecules is indicative of the kinetics of DNA replication if the BrdU densities decrease as replication proceeds. Peaks indicate replication initiation events, valleys indicate termination sites, positive gradients correspond to leftward replication forks, negative gradients to rightward forks, and sudden drops mark replication fork pause sites. B) *S. pombe* cell cycle synchronised cultures were grown in media with BrdU and harvested post S-phase at the indicated times. High-molecular weight DNA was then extracted, nanopore sequenced, and analysed. As an example, we show the BrdU detected on a single molecule from a culture treated with 2 µM BrdU and the corresponding population level median replication time from (7). In the BrdU plot, the grey dots indicate the probability of BrdU at each thymidine and the black line indicates the fraction of BrdU in 300 thymidine windows. In the replication time plot, time is shown increasing down the y-axis. In both plots the x-axis indicates the genomic position on chromosome III. C) Similarly to *S. pombe* cells, *S. cerevisiae* cells were grown in media with BrdU and harvested post S-phase at the indicated times. High-molecular weight DNA was then extracted, nanopore sequenced, and analysed. As an example, we show the BrdU detected in a single molecule from a culture treated with 30 µM BrdU and the corresponding population level median replication time from (21) plotted as in panel B. D) Cell cycle synchronised *S. pombe* were grown in the presence of 4 µM BrdU with a 400 µM thymidine 30 minutes after release. As an example, we show the BrdU detected on a single molecule and the corresponding population level median replication time plotted as in panel B. E) Fraction of BrdU incorporated over time in *S. cerevisiae* and *S. pombe* cells treated with a range of BrdU and thymidine concentrations. For each single molecule, the fraction of BrdU incorporated in every 1 kb window was determined and plotted in 5 min bins of population level median replication time from (21) and (7), for *S. cerevisiae* and *S. pombe*, respectively. The dotted black line indicates that only molecules with a BrdU fraction ≥0.05 are plotted. The black diamonds indicate the median BrdU fraction for each replication time interval.

In this study, we established the experimental conditions to perform DNAscent in *S. pombe* to map replication dynamics with high coverage and with single-molecule resolution. We identify thousands of molecules with replication fork direction, initiation, termination, and pause sites across the whole genome including the rDNA, the mating-type locus and the centromeres. We observe clear clustering of replication pause events at previously reported replication fork barriers in the rDNA and the mating-type locus. At the centromeres, we identify termination zones located within the central regions, initiation sites in the pericentromeric regions and an increase in pauses only in centromere 2. Overall, we show that DNAscent is a high-throughput method to study all major replication dynamic features— initiation, termination, fork direction and fork pauses—across the whole genome in fission yeast.

## MATERIAL AND METHODS

### Yeast strains

*Schizosaccharomyces pombe* (*S. pombe*) strain AW2224 (*h-, smt-0, leu1::leu1:Padh1-hENT1, gde1::his7:Padh1-hsvTK, cdc2-asM17, ade6-704*) was used in this study. Strain AW2224 was created by crossing YDP435 (*h-, smt-0, leu1::leu1:Padh1-hENT1, gde1::his7:Padh1-hsvTK, ade6-704*) and AW2181 (*h+, cdc2-asM17, ade6-704, leu1-32, ura4D18*) and random spore analysis was used to screen for the desired genotype (19). The *cdc2-asM17* allele was confirmed through sensitivity to growth on YE-agar media containing 3mM 3BrPP1. Intact *Padh1-hENT1* and *Padh1-hsvTK* constructs were confirmed using PCR.

*Saccharomyces cerevisiae* (*S. cerevisiae*) strain ARY017 (*MATa, RAD5, BUD4, leu2, ura3, trp1, ade2, his3 ChrVIII:202751::pSAC6-hENT1-tENO2-pPOP2-hsvTK-tTDH1*) was used in this study. To construct ARY017, the W303 strain T7107 was transformed with NotI digested pAR045 which contains *hENT1* and *hsvTK* which were codon optimised for expression in budding yeast.

### Cell cycle synchronisation, BrdU/thymidine treatment and flow cytometry

*S. pombe* cells were grown in YES medium (Formedium, PCM0310) at 30°C until reaching an OD_600_ = 0.3. Then, cells were arrested in the G2 phase by the addition of 3BrPP1 (stock concentration of 2 mM, Abcam, ab143756) at a 1:1000 dilution, followed by incubation for 3 hours. To resume cell cycle progression, 3BrPP1 was removed by vacuum filtration, and cells were resuspended in fresh YES medium at 30°C. Five minutes after filtration, BrdU (Sigma, B5002) was added to a final concentration of 0.5, 2, or 4 μM. For thymidine-treated samples, thymidine (Sigma, T9250) was added 30 minutes after media filtration at final concentrations of 40, 200, or 400 μM. Samples were harvested 60 minutes after media filtration, pelleted by centrifugation at 3,000 × g for 10 minutes, washed with PBS, and stored at -80°C until DNA extraction. Flow cytometry samples were collected before G2 arrest (asynchronous culture timepoint), immediately after release (0 min timepoint), and every 5 minutes thereafter until sample harvesting to monitor cell-cycle progression. Flow cytometry samples were treated with RNase A (Sigma, R6513) and proteinase K (Sigma, P2308) before DNA staining with SYTOX Green (Thermo Fisher, S7020), followed by sonication and analysis using a BD FACSAria Fusion cytometer. Flow cytometry profiles were analysed using FlowJo v10.8.1.

*S. cerevisiae* cells were grown in YPAD medium (Formedium, CCM1010) at 30°C until reaching OD_600_ = 0.3. Cells were arrested in the G1 phase by adding α-factor (Cambridge Bioscience, Y1001) to a final concentration of 0.5 μM. BrdU was added 95 minutes after α-factor addition at final concentrations of 10, 30, or 100 μM. Cell cycle progression was resumed 2 hours after α-factor addition by treating cells with pronase (Sigma, 53702) at 200 μg/ml. To prevent re-entry into a second cell cycle, nocodazole (Merck, 487928) was added at 15 μg/ml 20 minutes after pronase addition. Samples were harvested 90 minutes after pronase addition, pelleted by centrifugation at 3,000 × rpm for 5 minutes, washed with water, and stored at -80°C until DNA extraction. Flow cytometry samples were collected before G1 arrest (asynchronous culture timepoint), immediately before release (0 min timepoint), every 10 minutes thereafter for one hour, and before sample harvesting (90 min timepoint) to assess cell-cycle progression. Flow cytometry samples were processed and analysed as described for *S. pombe*.

Biological samples and their treatments are described in Supplementary Table S1.

### DNA extraction, library preparation and nanopore sequencing

High molecular weight DNA extraction from *S. pombe* and *S. cerevisiae* was performed using the Nanobind Tissue Kit (PacBio, 102-302-100) with modifications. This product is now the Nanobind PanDNA Kit (PacBio, 103-260-000). Briefly, 100 mg of cell pellets were resuspended in 1 ml of Spheroplasting Buffer 1 (100 mM Tris-HCl pH 9.5, 14 mM β-mercaptoethanol) followed by cell wall digestion by resuspending the cells in 400 µl of Spheroplasting Buffer 2 (1 M Sorbitol, 25 mM EDTA, 1.6 mM Citric Acid, 8.4 mM Sodium Citrate) and 100 µl of 50 mg/ml Zymolase 100 T (AMSBIO, 120493-1) in Spheroplasting Buffer 2. Cells were incubated for 1-1.5 h on a Thermomixer at 35 °C with 300 rpm shaking for 10 s every 5 min. Following digestion, spheroplasts were washed in PBS and resuspended in 20 μl Proteinase K, 50 μl Buffer SB, and 150 μl Buffer BL3. Samples were incubated at 55 °C in a ThermoMixer, shaking at 300 rpm for 90 minutes. Next, 20 μl of RNase was added, followed by an additional 90-minute incubation under the same conditions. Cells were centrifuged at 2,000 x g at room temperature for 3 min. The supernatant was transferred to a new tube using a wide-bore pipette and DNA precipitation and clean-up were performed according to the manufacturer’s instructions. For extraction of DNA from *S. cerevisiae*, Zymolase 20 T (AMSBIO, 120491-1) was used instead of Zymolase 100 T, and the RNase incubation step was reduced to 30 min. DNA concentration and integrity were assessed using the Qubit dsDNA HS assay kit (Invitrogen, Q33230) and TapeStation (Agilent Technologies).

Libraries were prepared using the ligation sequencing kit SQK-LSK109 (Oxford Nanopore Technologies; R9 chemistry) or SQK-LSK114 (Oxford Nanopore Technologies; R10 chemistry) as stated by the manufacturer. Libraries were loaded on MinION flow cells FLO-MIN106 (Oxford Nanopore Technologies; R9 chemistry) or FLO-MIN114 (Oxford Nanopore Technologies; R10 chemistry) as stated by the manufacturer.

Sequencing datasets are described in Supplementary Table S2.

### DNAscent pipeline

The detection of BrdU and the inference of replication fork direction, and initiation and termination sites, were performed using a DNAscent/FORKscent pipeline (pipeline_fast5_pod5_to_dnascent_dorado.sh) from https://github.com/DNAReplicationLab/fork_arrest. Briefly, nanopore basecalling was performed using Guppy version 5.0.7 (configuration file dna_r9.4.1_450bps_hac.cfg) or Guppy version 6.5.7 (configuration file dna_r10.4.1_e8.2_400bps_5khz_hac.cfg) for R9 or R10 data, respectively. Reads were aligned to references genome sequences using minimap2 version 2.24 (20). For *S. pombe* data, reads were aligned to either the reference genome ASM294v2 (https://www.ncbi.nlm.nih.gov/datasets/genome/GCF_000002945.1/), a custom rDNA array with 22 copies of rDNA, or a *de novo* assembly of the AW2224 strain. For *S. cerevisiae* data, reads were aligned to the reference genome sacCer3 (https://www.ncbi.nlm.nih.gov/datasets/genome/GCF_000146045.2/). For subsequent analyses only primary alignments with a read length ≥1,000 bp, were retained. Except for rDNA aligned data, only those alignments with a quality score ≥20 were retained. Alignment (bam files) and raw nanopore (fast5 files) data were then used to determine the probability of BrdU substitution at each thymidine position using DNAscent v2 or DNAscent v4 (https://github.com/MBoemo/DNAscent), for R9 or R10 data, respectively. BrdU probabilities were stored in the modbam file format to allow downstream analyses and visualisation with software tools like samtools, bedtools or modkit. To call replication fork direction, initiation and termination sites, we used the software forkSense from the DNAscent v2 package. ForkSense uses the BrdU probabilities to infer replication fork direction and initiation and termination sites.

### BrdU fraction over replication time

We compared the BrdU fraction on individual reads with the population median replication time. For *S. pombe*, the genome replication timing profile data, in 1 kb windows and in 1 minute intervals, was obtained from (7), accession number GSE62108. For *S. cerevisiae*, the genome replication timing profile data, in 1 kb windows and in 1 minute intervals, was obtained from (21), accession number GSE48212. The *S. cerevisiae* data was lifted from the sacCer1 to the sacCer3 genome assemblies using liftOver (22). For each 1 kb replication timing window, the fraction of BrdU (mean BrdU density using a probability threshold of 0.5) was determined for each read that overlapped for ≥100 thymidines. Levels of BrdU substitution with respect to replication time were then analysed in 5 minute intervals and visualised as violin plots.

### BrdU probability and fraction over genomic coordinates in single molecules

To plot the BrdU probabilities and fractions (or levels of BrdU substitution) from single molecules, we first extracted the modification probabilities from the mod.bam file containing the read of interest. Then, the level of BrdU substitution was determined in 300-thymidine windows using a BrdU probability threshold of ≥0.5. The modification probability and the BrdU substitution data for a single read were then plotted relative to genomic coordinates. BrdU probabilities and substitution level were plotted as grey dots and a black line, respectively.

### Heatmap of BrdU fractions

To produce a heatmap of BrdU substitution level along multiple reads, we extracted the modification probabilities using modkit (https://github.com/nanoporetech/modkit). Then, we applied a threshold of 0.5, so that modification probabilities <0.5 were converted to 0 and probabilities ≥0.5 were converted to 1. These values were averaged, in windows of 1000 nucleotides, to determine the fraction of BrdU substitution.

### Fraction of leftward-moving forks

To calculate the fraction of leftward replication forks, we counted the number of left and right forks at every base using bedtools genomecov, averaged the counts per 1000 bins and calculated the fraction of left forks by dividing the number of left forks by the total number of forks. At each genomic interval, we compared the fraction of leftward forks from all nascent single molecules to the equivalent value from Pu-seq data. To assess correlation, we calculated the Pearson correlation coefficient. The Pu-seq fraction leftward replication forks data (GSE62108_PU-seq_leftward_moving_fork.wig) was obtained from (7).

### Replication initiation and termination site count

To visualise DNAscent initiation and termination site frequency across the genome, we first calculated the midpoint of each DNAscent initiation/termination site. Then, the number of initiation/termination site midpoints was determined in 2 kb windows across the genome.

### Comparison of DNAscent replication initiation sites to published datasets

To calculate the closest distance from the DNAscent initiation sites to initiation sites from independent published datasets, we used bedtools closest. Then, we grouped the distances in 1 kb bins, counted the number of events per bin, and plotted the count as a histogram in 1 kb bins. DNAscent initiation sites from the mitochondrial genome were excluded from the analysis. To evaluate the significance of the distribution of distances, we compared them to the average distance from 1000 independent randomisations of the DNAscent initiation sites. For each randomisation, the observed DNAscent initiation site genomic intervals were shuffled (bedtools shuffle) across the reference genome.

We compare previously reported (Pu-seq) initiation site efficiencies with the number of observed DNAscent initiation events. For each Pu-seq initiation site we determined the number of DNAscent initiation site midpoints within 1 kb (either side). We then plotted the DNAscent initiation site counts with respect to the Pu-seq initiation efficiency as jitter dots with boxplots.

The file with Pu-seq initiation sites (GSE62108_origin-effici.bedgraph) was obtained from (7). Initiation sites from (5, 23, 24) where obtained from OriDB (25).

### Comparison of DNAscent and Pu-seq replication termination sites

To compare the DNAscent termination site midpoint counts with the Pu-seq termination event frequencies, we used 1 kb windows (sliding by 300 bp) since this is how the Pu-seq termination data was reported. The data were analysed and plotted in bins of 2% Pu-seq termination frequency. The file with Pu-seq termination frequencies (GSE62108_PU-seq_terminaton-events.wig) was obtained from (7).

### Pause site detection

To detect pause sites on single molecules, we used the rDNA_detect.py pipeline from https://github.com/DNAReplicationLab/fork_arrest. Briefly, we selected those molecules with a mean BrdU fraction of at least 0.05 and measured the BrdU fraction difference between an upstream and downstream window at each thymidine per molecule (window size 3x 290T or approximately 3 kb in the yeast genome). Then, we detected peaks in the density difference, requiring that peaks are at least 15x 290T apart on the genome (approximately 15 kb). Then, we collated peaks in density difference across all molecules, retaining only those with step sizes greater than 3x the standard deviation of all step sizes across all molecules.

We filtered out those peaks which were closer than 5 kb to the ends of molecules, and those with maximum and minimum BrdU densities ≤0.5 or ≤0.01, respectively, in their upstream or downstream windows.

### Visualisation of replication dynamics

To visualise the location of left and right forks, initiation sites, termination sites and pause sites, we used custom R scripts that took the genomic coordinates of each feature in the reads of interest, and represented them as blue arrows (left forks), right arrows (right forks), black rectangles (initiation sites), grey rectangles (termination sites), green triangles (pause sites in leading strand) and pink triangles (pause sites in lagging strand).

To visualise the fraction of leftward-moving forks, initiation event counts, origin efficiencies, termination event counts, termination event frequencies and pause event counts we used DnaFeaturesViewer (https://edinburgh-genome-foundry.github.io/DnaFeaturesViewer/).

### Design of the rDNA sequence

To study the ribosomal DNA (rDNA), we generated an rDNA sequence with 22 copies of the rDNA unit with the replication fork barrier (RFB) region in the middle. Each rDNA unit consists of a 10.9 kb sequence obtained from the reference genome ASM294v2, coordinates chrIII:5539-16411. The rDNA sequence was annotated using the gene coordinates from the reference genome ASM294v2, the origin of replication ARS3001 coordinates from (26) (GenBank: AF040270.1) and the locations of the RFB based on (27, 28).

### *De novo* assembly

To obtain a reference sequence for strain AW2224, we used high-quality long reads from the sequencing dataset IDS_65-R9 (see Supplementary Table S2). Only reads with a quality score >9 and read length ≥75 kb were used. We then performed a *de novo* assembly using Flye (v2.9) with an estimated genome size of 14 Mb. The resulting contigs were evaluated by comparing them to the previously published assembly (29) and the reference genome ASM294v2 using MUMmer (v3.23). All *S. pombe* chromosomes were recovered as single contigs, and all contigs had a sequencing coverage >30.

### Annotation of the mating-type locus and the centromeres

To annotate the mating-type locus in the reference sequence for strain AW2224, *mat* genes and flanking genes were identified using BLAST+ (v2.9.0) (30), with known gene sequences from ASM294v2 as queries. The coordinates of the replication fork barrier *RTS1* were determined by aligning its partial sequence from (31), while those of replication fork barrier *MPS1* were identified using the *MPS1* sequence from (32). The 263-bp *smt-0* deletion (33) was confirmed by aligning the mating-type locus from strain AW2224 with the mating-type locus from ASM294v2.

Centromeric repeats, core elements and flanking genes were identified using BLAST+ (v2.9.0), with known repeat, core and flanking gene sequences from ASM294v2 as queries. The locations of tRNA genes were predicted using tRNAscan-SE (34).

## RESULTS

### BrdU incorporation in fission yeast serves as a proxy for replication time

Nanopore sequencing enables inference of DNA replication dynamics through the detection of synthetic analogues of thymidine, such as bromodeoxyuridine (BrdU), that are incorporated into nascent strands during S-phase in a time-dependent manner (Figure 1A) (12, 13). This method was established in *S. cerevisiae*, where BrdU levels incorporated into the DNA naturally decrease as replication progresses (12, 35). To detect BrdU incorporation, cells are fed BrdU, harvested post-S phase, high-molecular weight genomic DNA extracted and subjected to nanopore sequencing. The resulting sequence data are basecalled, aligned to the reference genome sequence and sites of BrdU incorporation are determined using the DNAscent software (see Methods). The DNAscent software assigns a probability of BrdU substitution (grey circles; Figure 1A) to each reference thymidine position across every sequencing read. Positions with a BrdU probability ≥0.5 are considered positive BrdU calls, and the fraction of BrdU incorporation is calculated in 300-thymidine windows across each sequencing read (black lines; Figure 1A).

To assess the profiles of BrdU incorporation in *S. pombe* and *S. cerevisiae*, we treated synchronised cells with a range of BrdU concentrations during a single S phase. None of the BrdU treatments affected cell cycle progression, as shown by flow cytometry profiles of cellular DNA content (Supplementary Figure 1). In *S. pombe*, we observed that at the single-molecule level, BrdU incorporation was constant along the DNA and did not correlate with previously published median replication time from a cell population (Figure 1B). In contrast, *S. cerevisiae* cells exhibited BrdU incorporation gradients that correlated with cell population median replication timing data (Figure 1C). To obtain BrdU gradients in *S. pombe*, we added a thymidine chase 30 minutes after the BrdU treatment – a time that corresponds with mid-S phase (Figure 1D and Supplementary Figure 1). In the same genomic location as shown in Figure 1B, we observed that the single molecules had BrdU incorporation levels resembling the median replication time. This result indicated that a thymidine chase led to BrdU decrease with respect to replication time. To further assess the relationship between BrdU incorporation and replication time, we plotted all single-molecule BrdU fractions against population level median replication times. In *S. cerevisiae*, BrdU incorporation decreased with later replication times at all tested concentrations (Figure 1E; top), supporting the use of BrdU levels as a replication timing proxy. These observations are consistent with previous reports from our group and others (12, 35). In contrast, *S. pombe* exhibited no decline in BrdU incorporation with replication time unless a thymidine chase was applied (Figure 1E; middle and bottom). Equivalent results were obtained using the newest nanopore sequencing chemistry (R10), although we observe that BrdU detection in R10 data was noisier, especially at late replication times (Supplementary Figure 2). These results indicate that BrdU incorporation kinetics differ between *S. cerevisiae* and *S. pombe*, and that in *S. pombe*, replication timing cannot be inferred from endogenous BrdU incorporation patterns alone. However, a thymidine chase produces BrdU gradients that correlate with replication time, indicating that single-molecule BrdU fractions can also serve as a proxy of replication time in *S. pombe*.

### BrdU incorporation levels indicate replication fork direction, initiation and termination sites

In *S. pombe*, the addition of a thymidine chase in mid-S-phase resulted in high levels of BrdU incorporation followed by gradients of declining BrdU levels as replication proceeded (Figure 1D). These gradients resemble those observed in *S. cerevisiae* (12, 13) (Figure 1C) which allowed us to computationally infer replication fork direction, and call initiation and termination sites on individual sequencing reads using the forkSense software (13). Positive BrdU gradients are identified as leftward-moving forks (blue arrows), whereas negative BrdU gradients are identified as rightward-moving forks (red arrows) (Figure 2A). Diverging leftward- and rightward-moving forks indicate initiation sites (black rectangles) and converging leftward- and rightward-moving forks indicate termination sites (grey rectangles). In a dataset from *S. pombe* cells treated with 4 µM BrdU and 400 µM thymidine, we obtained 462,052 nascent single molecules in which we identified 370,298 forks, 65,098 initiation sites and 39,157 termination sites. As an example, we show all the single molecules with ≥1 forks in a 100-kb region from chromosome I (Figure 2B and C). BrdU densities are represented as a heatmap in Figure 2B with one single molecule per row. The aligned single molecules clearly show variable levels of BrdU incorporation across this genomic locus. Most single molecule reads show a peak of high BrdU incorporation (green-yellow; Figure 2B) around position 845 kb indicating early replication, in contrast to a region with low BrdU incorporation (blue; Figure 2B) at around 865 kb indicating later replication. Declining levels of BrdU incorporation to the left and right indicate progressively later replication times and therefore replication fork direction. Transitions from left- to rightward forks are identified as initiation sites and right- to leftward as termination sites. Across the example locus on chromosome I, we observe alternating regions of predominantly leftward (blue arrows) and rightward (red arrows) fork direction (Figure 2C). The position of transitions from leftward to rightward fork direction are similar between single molecules, for example at ∼845 kb where most molecules have diverging forks indicating an initiation site. By contrast, there is more variability between molecules in the position of converging forks (for example at ∼865 kb) that indicate termination sites.

**Figure 2:**
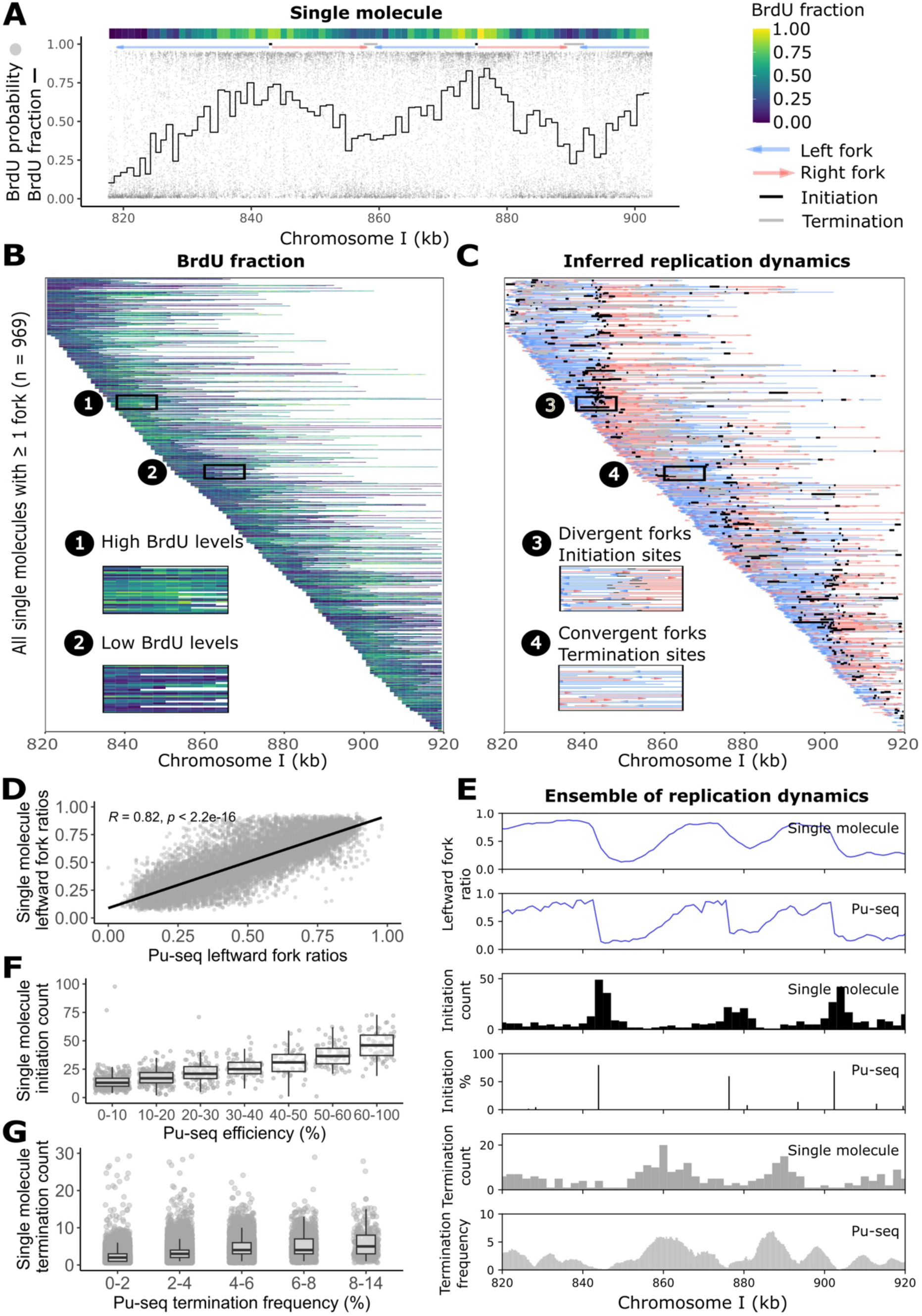
BrdU incorporation levels reveal replication fork direction, location and efficiency of initiation sites and location of termination sites. Single molecules were aligned to the reference genome ASM294v2. A) Example of a nascent single molecule read with inferred replication dynamics. BrdU probability and fraction are plotted against genomic coordinates (as in Fig. 1B). The lines above the BrdU probabilities indicate the position of right-moving forks (red arrow), left-moving forks (blue arrow), initiation sites (black rectangle) and termination sites (grey rectangle). The heatmap indicates the BrdU fraction. B) Heatmap of BrdU fractions in single molecules with ≥1 fork. Two example areas (1 and 2) are shown zoomed in and highlighted in black. C) Replication fork directions, initiations and terminations on all single molecules from panel B. Features are represented as in the example single molecule from panel A. As in panel B, two example areas (3 and 4) are zoomed in and highlighted in black rectangles. D) Fraction of leftward-moving forks identified on single molecules and Pu-seq from (1). E) Comparison of replication dynamics from single molecule sequencing and Pu-seq at the example locus. The panels show, from top to bottom: fraction of leftward-moving forks detected on single molecules, fraction of leftward-moving forks detected with Pu-seq, count of single molecules initiation sites, efficiency of Pu-seq origins, count of single molecules termination sites, and frequency of termination events detected with Pu-seq. Pu-seq data from (1). F) Single-molecule initiation counts versus Pu-seq origin efficiencies. The count of single-molecule initiation site midpoints within +/- 1kb from a Pu-seq origin were grouped by Pu-seq efficiency. Only DNAscent initiation counts ≤60 are shown. G) Single-molecule termination counts versus Pu-seq termination frequencies. Only DNAscent termination counts ≤30 are shown.

To validate the single molecule replication fork direction, and initiation and termination sites, we compared them to those identified by previous cell-population studies (5, 7, 23, 24). To allow direct comparison with the cell population data from these studies, we calculated an ensemble of our single-molecule data. For replication fork direction, we calculated the fraction of leftward-moving forks (1 kb window, 300 bp steps) across the genome using all our single-molecule data. For initiation and termination sites, we counted the number of events in 2 kb windows. When comparing the ensemble single molecule and Pu-seq fraction of leftward replication forks, we observed a strong correlation (*R* = 0.82) between the datasets (Figure 2D). At the example genomic locus, we see close agreement between the ensemble fraction of leftward forks from our single-molecule data and the Pu-seq data (Figure 2E). We then compared the single-molecule initiation and termination sites, to those inferred from Pu-seq data. Half of the single-molecule initiation events (33,428 out of 60,731 sites) were within 2 kb of a Pu-seq initiation site, and this intersection was highly statistically significant (Supplementary Figure 3A). Similar results were obtained when looking at the intersection of single-molecule initiation sites and other independent studies (Supplementary Figure 3B-D). The Pu-seq methodology determines both the location and efficiency of origins (fraction of cells in which the origin initiates DNA replication). Our dataset has a ∼100x coverage of reads with ≥1 initiation site, therefore, we reasoned that the number of times an initiation was observed on single molecules would indicate the efficiency of the origin. To test this, we compared the count of single-molecule initiations with the Pu-seq efficiency measurement at each Pu-seq origins (Figure 2F). We observe a positive correlation between the count of single-molecule initiation events and Pu-seq origin efficiency. We also found a positive correlation between the count of single-molecule termination events and frequency of Pu-seq terminations (Figure 2G). This is reflected in the example locus (Figure 2E), where there is a striking similarity between the single molecule and the Pu-seq data for fraction of leftward-moving forks, and location and efficiency of initiation and termination sites. Together these findings validate that the DNAscent methodology provides single molecule information on replication fork direction and the location of initiation and termination events. In addition, ensembles of single molecules provide a measure of the frequency of replication initiation and termination across the genome.

### BrdU incorporation reveals replication pause locations in leading and lagging strands at the rDNA replication fork barrier

We have shown that BrdU incorporation levels in single DNA molecules can be used to infer replication fork directions, as well as initiation and termination sites. BrdU densities can also indicate the location of replication pauses, as previously demonstrated in *S. cerevisiae* (12, 14) where sudden drops in BrdU density denote replication pauses within the ribosomal DNA (rDNA). Fission yeast rDNA consists of 100–150 tandem repeats, each containing the ribosomal RNA genes, a replication origin, and a region containing four replication fork barriers (RFBs) (Figure 3A) (36). RFBs are located downstream of the largest ribosomal gene to arrest rightward-moving forks, thereby limiting replication in the opposite direction to transcription (37). Thus, we anticipated that we would observe a sudden drop in BrdU density indicating replication pauses at the RFB. First, we aligned our single-molecule dataset (4 µM BrdU / 400 µM thymidine) to a reference sequence of 22 rDNA repeats. Second, we used DNAscent to detect BrdU and forkSense to infer the position of replication forks, initiation and termination events. Third, paused replication forks were computationally identified from large decreases in BrdU density (see Methods), thus prioritising a low false positives rate at the expense of identifying shorter pauses that would present as a smaller decrease in BrdU density (14). Among the 8,165 nascent reads that aligned to rDNA, we identified 12,677 forks, 3,654 initiation sites, 2,079 termination sites, and 376 pauses. Finally, combining information about fork direction and sequenced strand, we labelled each pause site as from a leading or lagging strands.

**Figure 3.**
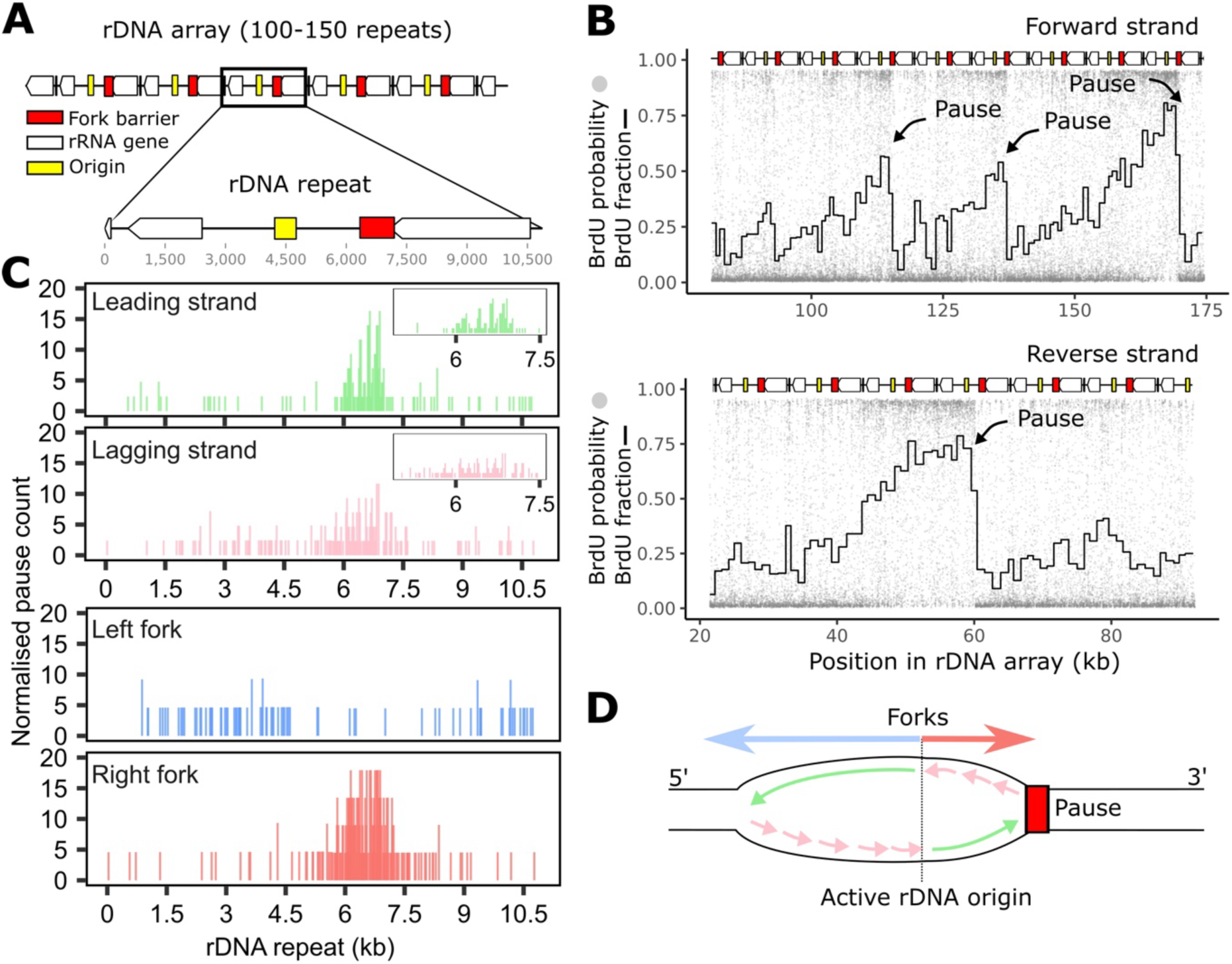
DNA replication dynamics at the *S. pombe* rDNA. Single molecules were aligned to a custom rDNA array containing 22 rDNA repeats. A) Schematic of the *S. pombe* rDNA. Each repeat contains rRNA genes, a replication origin (ARS3001), and a region with fork barriers. B) Example of two single molecules that align to the rDNA. A sharp drop in BrdU incorporation indicates the location of a pause in replication. BrdU probability and fraction are plotted against the custom rDNA array coordinates. The rDNA array is represented on top of the molecules. C) Normalised pause count per rDNA unit. Pauses were collapsed in one rDNA unit and separated depending on whether they were found in the lagging/leading strand or left/right fork. D) Schematic of replication at the rDNA. After an origin fires, the left forks progresses while the right fork is paused at the barriers on both the leading and lagging strand.

In example single molecules, aligned to the custom rDNA sequence, we observe clear drops in BrdU density that align with RFBs (Figure 3B). Interestingly, these molecules show that only a fraction of the origins of replication were active (∼1 in 3), consistent with observations from DNA combing (10). Then, we visualised all reads containing ≥1 pause and observed that anticipated drops in BrdU density aligned to the RFB (Supplementary Figure 4A). We also observed that most initiation sites overlapped with the previously characterised replication origin, and that rDNA units are predominantly replicated by leftward forks, thus ensuring co-directionality of transcription and replication (Supplementary Figure 4B). Next, we collapsed the replication pause site data onto a single representative rDNA repeat. The identified sites of pausing were predominantly on rightward-moving forks and enriched at the RFB location (Figure 3C). We observe that the leading strand pauses are more tightly clustered and closer to the RFB than the lagging strand pauses. These results confirm that the DNAscent methodology can identify replication pause sites and reveal the strand of synthesis and fork direction, giving a more detailed and quantitative view of replication dynamics at the rDNA (Figure 3D).

### Frequent replication pausing in the mating-type locus

The ability to detect pauses in replication within the rDNA prompted us to investigate pauses throughout the unique genome. To test this, we mapped the data to a *de novo* assembly of our strain AW2224 (see Methods). This allowed us to analyse genomic regions that differ between this strain and the reference genome, including the centromeres and mating-type locus. Across the whole unique genome, the most prominent identified replication pause site was within the mating-type locus (Supplementary Figure 5). The mating type of *S. pombe* is determined by the *mat1* locus located at chromosome II. Flanking the *mat1* gene, there are two elements that control the direction of replication across the *mat1* locus: *RTS1* and *MPS1* (38) (Figure 4A). The *RTS1* element is downstream of *mat1* and acts as a polar terminator of replication, terminating rightward-moving forks (39). The *MPS1* element is upstream of *mat1* and acts as a polar pause, stalling leftward-moving forks (40). Thus, the *mat1* locus is predominantly replicated by leftward-moving forks that occasionally pause at the *MPS1* site. This tightly controlled replication pattern is proposed to allow the formation of an imprint (ribonucleotides and/or a nick) at the *MPS1* site, which can lead to mating-type switching (41). However, the frequency of the pausing at this site *in vivo* is not known.

**Figure 4.**
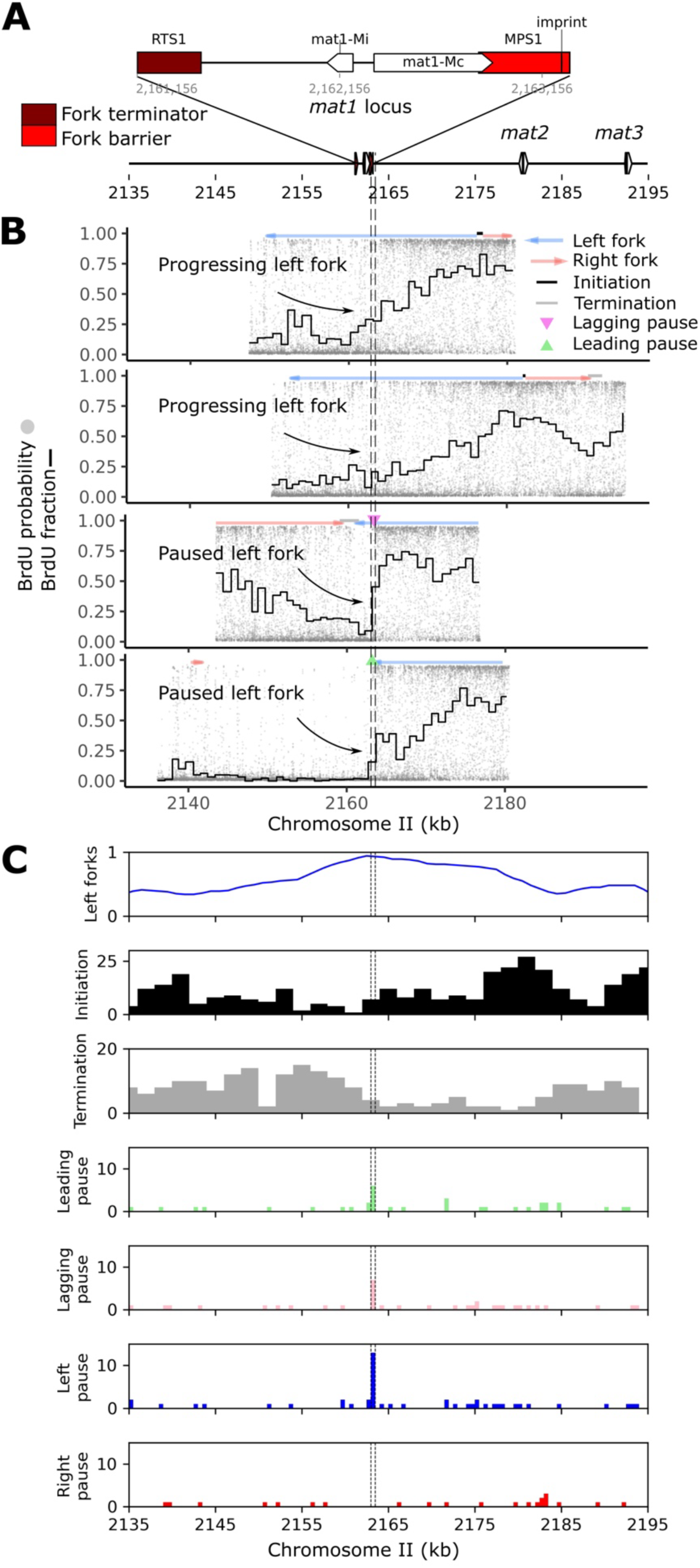
DNA replication dynamics at the *S. pombe* mating-type locus. Single molecules were aligned to a *de novo* whole-genome assembly from strain AW2224. A) Schematic overview of the mating-type locus in *S. pombe*. The mating-type locus is comprised of three genes: *mat1*, *mat2* and *mat3* in chromosome II. At the *mat1* locus, the replication terminator *RTS1* prevents rightward-moving forks across the *mat1* genes, while the replication fork barrier *MPS1* occasionally pauses leftward-moving forks. B) Examples of single molecules showing unperturbed left fork progression or stalled left fork at the *MPS1* site. BrdU probability and fraction are plotted on the coordinates for *de novo* genome assembly and visualised as in Fig. 1B. C) Ensemble of single-molecule replication dynamics at the mating-type locus. The panels show from top to bottom: fraction of leftward-moving forks, count of initiation sites, count of termination sites, count of pause sites in the leading strand, count of pause sites in the lagging strand, count of pause sites in left forks, and count of pause sites in right forks. Single-molecule initiation and termination were counted in 2 kb genomic windows and pause sites were counted in 500 bp windows.

Our single-molecule dataset allowed us to observe two distinct patterns of replication at the *mat1* locus: continuous replication by a leftward fork, or leftward-moving fork pauses at the *MPS1* site (Figure 4B). Unperturbed fork progression was the most frequent scenario (∼60 % of molecules) (data not shown). Ensemble of replication features showed a clear termination area upstream *mat1* which aligns with the expectation of right forks terminated at the *RTS1* element (Figure 4C). All pauses were found in left forks, and mapped to the *MPS1* element. These results indicate that the *mat1* locus is almost exclusively replicated by a leftward-moving fork (94%) that may pause (∼40% of molecules) at the *MPS1* site or continue through *mat1*, and that most rightward-moving forks terminate at the *RTS1* site.

### Replication dynamics across the highly repetitive centromeres

The high levels of repetitive sequence across *S. pombe* centromeres have precluded comprehensive analysis of the replication dynamics by previous short-read based genomic studies. Each of the three centromeres is comprised of a central core flanked by inner (*imr*) and outer (*dg* and *dh*) repeats containing tRNAs (Figure 5A) (42). The inner repeats are unique for each centromere, but the outer repeat sequences are shared among centromeres and are heterochromatic. The central core is the site of kinetochore assembly; the sequence of centromere 2 central core (*cnt2*) is unique, while those of centromeres 1 and 3 (*cnt1* and *cnt3*) are highly similar. Origins of replication have been found in the repeats of centromere 2 but not within the central core (43, 44), and it has been proposed that the centromeric tRNAs act as barriers to DNA replication (45). Therefore, we sought to use the single-molecule replication dynamics dataset to comprehensively analyse the landscape of replication across all three *S. pombe* centromeres.

**Figure 5.**
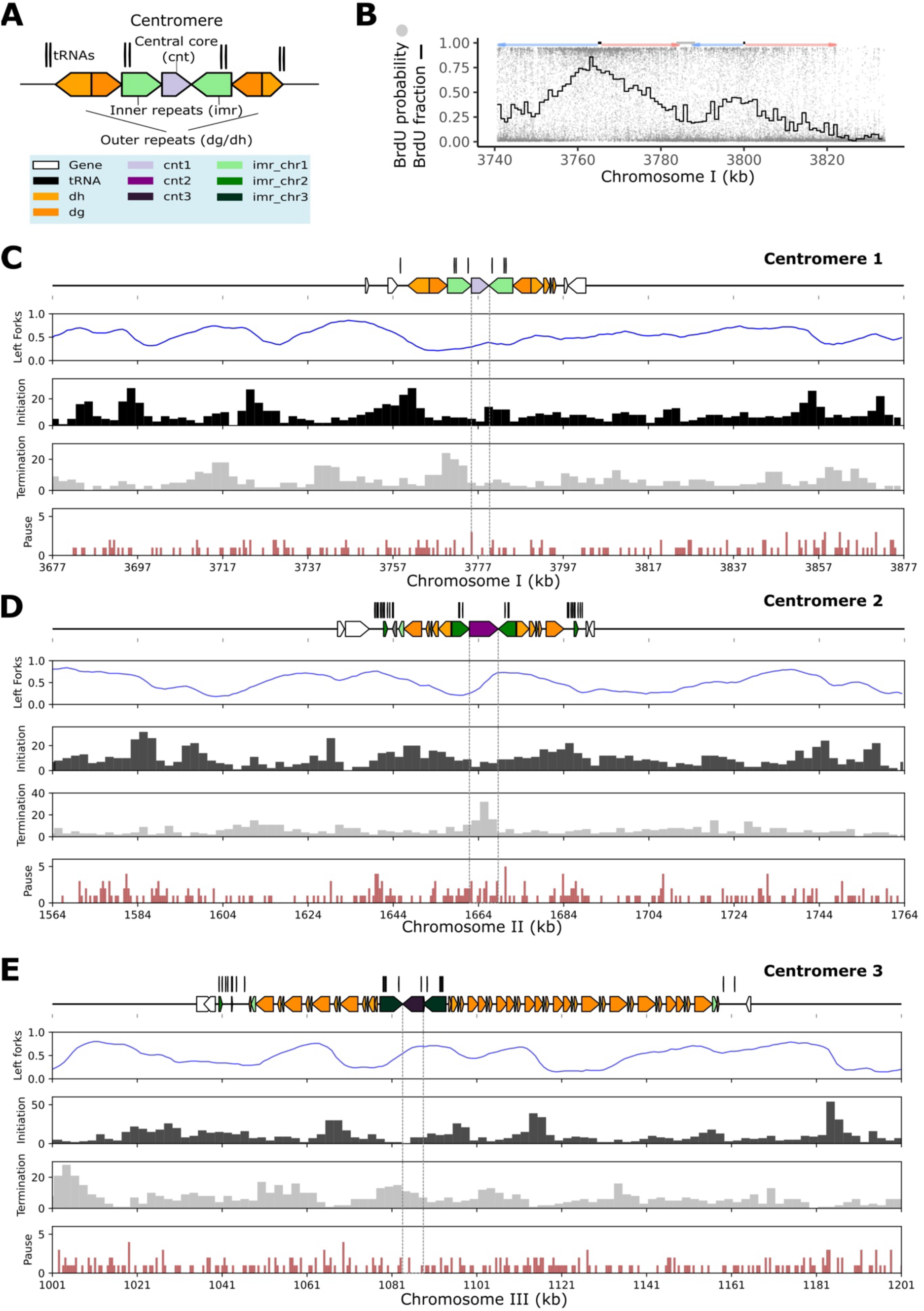
DNA replication dynamics at the *S. pombe* centromeres. Single molecules were aligned to a *de novo* whole-genome assembly from strain AW2224. A) *S. pombe* centromeres are comprised of outer repeats (*dg* and *dh*), flanking a central region composed of inner repeats (*imr*) and a central core (*cnt*). tRNAs are found within and adjacent to each centromere. B) Examples of a single molecule that aligns to centromere 1 plotted on the coordinates for *de novo* genome assembly and visualised as in Fig. 1B. C) Ensemble of single-molecule replication dynamics across centromere 1. The panels show from top to bottom: centromere annotation track with the tRNAs on a separate y-axis (for clarity only two flanking genes are shown either side of the centromere), fraction of leftward forks, count of initiation sites, count of termination sites and the count of pause sites. Single-molecule initiation and termination sites were counted in 2 kb genomic windows whereas the pause sites were counted in 500 bp windows. The central cores are highlighted with dashed lines. D) Ensemble of single-molecule replication dynamics across centromere 2 plotted as in panel C. E) Ensemble of single-molecule replication dynamics across centromere 3 plotted as in panel C.

After careful examination of published *S. pombe* genomes sequences we determined that the centromeric sequences were either incomplete or showed significant structural differences compared to our data (not shown). Therefore, we used our *de novo* genome assembly and annotated the centromeric regions for our strain (see Methods). Using this assembly as a reference for DNAscent and forkSense analyses, we identified 4,748 forks, 1,116 initiations, 729 terminations, and 285 pauses intersecting with the three centromeres. As an example, we show a single molecule that aligned to centromere 1 (Figure 5B). Across centromere 1, most initiation sites were located outside the central core, particularly in the first *dh* repeat (Figure 5C). Termination sites were associated with the inner repeat downstream of the central core, and replication pauses were distributed throughout the centromere. For centromere 2, initiation sites were found throughout the centromere, especially in the outer repeats (Figure 5D). A distinct enrichment in termination sites was observed across the central core, and notably, replication pauses were most frequent in proximity to the tRNAs within the outer repeat sequences. Centromere 3 DNA replication resembled centromere 1, with clustered initiation sites in the outer repeats and termination sites both in the central domain and in the repeats, but with no clear enrichment of pauses (Figure 5E). These findings indicate that all centromeres have replication origins outside the central core, termination events within the central region, and that centromere 2 may be unique in showing replication pausing close to tRNAs.

## DISCUSSION

In this study, we demonstrated that direct nanopore sequencing and detection of the nucleotide analogue BrdU can determine replication dynamics across the *S. pombe* genome. This methodology, which we named DNAscent, captures all major features of DNA replication—initiation, fork progression, termination, and pausing—on single molecules.

Additionally, it has the advantage of reporting the strand of synthesis which allows pauses on leading and lagging strands to be distinguished. Moreover, nanopore sequencing enabled >200× coverage of long (>30 kb, up to ∼120 kb) nascent molecules genome-wide in a single MinION run. Such high coverage facilitates detection of rare events that are challenging to resolve with population-level methods, and also makes it possible to detect replication dynamics in repetitive sequences that cannot be studied with short-read sequencing methods. To validate our findings, we compared them against several independent published datasets (Figure 2 and Supplementary Figure 3). Initiation sites aligned with those identified in prior studies, with higher counts correlating with more efficient initiation sites. Fork direction and termination sites also correlated with published Pu-seq data (7). Finally, most frequent pauses in the rDNA and the rest of the genome correlated with well-characterised replication fork barriers (Figure 3 and 4). Altogether, this approach provides a detailed view of replication dynamics across the whole genome that complements and extends previous studies (5–7, 10, 11, 23, 24).

To map replication dynamics, we measured changes in BrdU concentration with respect to time. In *S. cerevisiae*, endogenous mechanisms lead to decreases in BrdU levels over time, whereas in *S. pombe* BrdU levels were constant throughout S-phase (Figure 1). Therefore, in this regard *S. pombe* is like other eukaryotes (15) and suggests different mechanisms of dNTPs pool regulation in budding yeast compared to other eukaryotes. Although dTTP levels have been reported to rise in fission yeast during S-phase (46), the magnitude of this increase is notably less than in budding yeast. This may reflect species-specific regulation of ribonucleotide reductase, the enzyme responsible for deoxyribonucleotide biosynthesis (47). Nonetheless, we demonstrate that in fission yeast a thymidine chase can be used to experimentally reduce BrdU levels with respect to time. Similar approaches might be used for other organisms that, like fission yeast, do not naturally show BrdU gradients.

The repetitive rDNA locus, comprising 100–150 tandem 10.9 kb units, poses particular challenges for replication analysis. Our long-read sequencing approach allowed us to resolve replication patterns across multiple contiguous rDNA repeats, revealing two primary modes: active initiation within the rDNA repeat (∼1 active origin in 3 rDNA repeats), and passive replication by forks entering from upstream (Figure 3B). Replication fork barriers shaped these patterns, and we observed a concentration of replication pausing events near a region with four replication fork barriers (48). Notably, lagging-strand pauses showed greater positional variability than those on the leading strand, consistent with the discontinuous synthesis of Okazaki fragments. Leading-strand pauses, by contrast, often occurred at four distinct positions that likely correspond to the four replication barriers (Figure 3C).

A *de novo* assembly allowed us to look at single-molecule replication dynamics in under studied areas, such as the mating-type locus and the centromeres. The mating-type locus is comprised of a heterochromatic silenced region with the *mat2* and *mat3* genes, and an actively transcribed region with *mat1* flanked by a replication termination site (*RTS1*) and a replication pause site (*MPS1*). We observed replication fork pausing at the *MPS1* site, leading to leftward replication and a downstream shift in termination (Figure 4). While this behaviour aligns with previous models (31, 32), it is now directly visualised at the single-molecule level for the first time and it allowed us to estimate that ∼40% of the cells pause in this location. In all centromeres, we found replication initiation sites within the outer centromeric repeats, whereas the central region functions as a termination zone (Figure 5). These findings are consistent with models proposing that replication dynamics are spatially organised to preserve centromere function (49). We also observed a modest enrichment of replication pausing at tRNA genes within centromere 2. This includes the tRNA^Ala^ gene in the *imr2* repeat, previously identified as a chromatin barrier (50). To our knowledge, this is the first direct observation of increased fork pausing at this locus, linking replication dynamics to known chromatin features in centromeric regions.

We also identified parallels between the organisation of the mating-type locus and centromeres. The *mat2* and *mat3* region shares heterochromatic features with the outer centromeric repeats, while *mat1*, like the centromere central cores, is euchromatic and associated with late replication. These similarities suggest a conserved strategy in which replication timing and pausing contribute to local chromatin structure. Fork pausing at the *MPS1* site may facilitate lagging-strand processes such as primer placement or chromatin modification (38). A comparable mechanism has been proposed for centromere 2, where pausing near flanking tRNA genes appears to protect the central core from inappropriate histone deposition (45). Interestingly, such pausing was absent at centromeres 1 and 3, indicating a different type of regulation.

In the future, DNAscent, alongside other nanopore-based methods such as FORK-seq (51), offer new opportunities for studying replication at single-molecule resolution. In particular, the ability to calculate fork velocity across different chromatin states will allow testing of whether replication is slowed in heterochromatin. Fission yeast offers a powerful model for this, given its compact genome and presence of mammalian-like heterochromatin, which is absent in budding yeast (52). Furthermore, the capacity to track multiple molecular signals— including DNA modifications like methylation—on the same single molecule paves the way for integrated analyses of epigenetic and replication dynamics (53). Future studies could systematically map replisome pausing sites (14) and extend these analyses to mutants affecting chromatin structure (e.g. *sir2*, *clr3*), chromosome organisation (e.g. *swi6*), or replication fork stability (e.g. *swi1*, *swi3*).

In summary, our findings establish DNAscent as a powerful tool for dissecting replication dynamics at single-molecule resolution in *S. pombe*. These principles hold true in more complex eukaryotes and can be applied to any organism that can incorporate BrdU.

## DATA AVAILABILITY

BrdU calls on all aligned reads (mod.bam format), inferred replication dynamics (bedgraph files), reference sequences, annotations and replication timing datasets are available from Zenodo under the DOI 10.5281/zenodo.15365203. The deposited files are listed in Supplementary Table S3. Code described here is available from the GitHub repositories (https://github.com/DNAReplicationLab/pombe_replication and https://github.com/DNAReplicationLab/fork_arrest).

Data will be publicly available after the review process. Please contact the corresponding author for access before then. Scripts were written in shell script, python, and R.

## SUPPLEMENTARY DATA

Supplementary Data are available online.

## AUTHOR CONTRIBUTIONS

Isabel Díez-Santos: Conceptualization, Formal analysis, Investigation, Methodology, Visualization, Writing – original draft, Writing – review & editing. Sathish Thiyagarajan: Methodology, Software, Writing – review & editing. Anna M. Rogers: Investigation, Writing – review & editing. Adam T. Watson: Methodology. Antony M. Carr: Funding acquisition, Resources, Supervision, Writing – review & editing. Conrad A. Nieduszynski: Conceptualization, Funding acquisition, Resources, Supervision, Writing – original draft, Writing – review & editing.

## ACKNOWLEDGEMENTS

We are grateful to colleagues at the Earlham Institute for their support and expertise: Dr. Andrew Goldson for guidance on flow cytometry, Grant Bexson for laboratory assistance, and Dr. Darren Heavens for advice on library preparation. We also extend our thanks to Dr. Martin Ayling, Sam Gallop, and Kamil Hepak from the Norwich Biosciences Institutes Research Computing team for their assistance with software installation and maintenance. We acknowledge Dr. Emma Heron and Dr. Yasukazu Daigaku for their prior work on *S. pombe* and for providing the YDP435 strain. The T7107 strain was kindly gifted by Prof. Tomoyuki Tanaka. Finally, we sincerely thank Prof. Crisanto Gutiérrez, Dr. Bénédicte Desvoyes, and Nerea Murugarren for their critical reading of the manuscript and valuable feedback.

