## Supplementary figures and tables for "Single-molecule sequencing maps replication dynamics across the fission yeast genome, including centromeres"

**SUPPLEMENTARY DATA**

**Supplementary Figure 1**

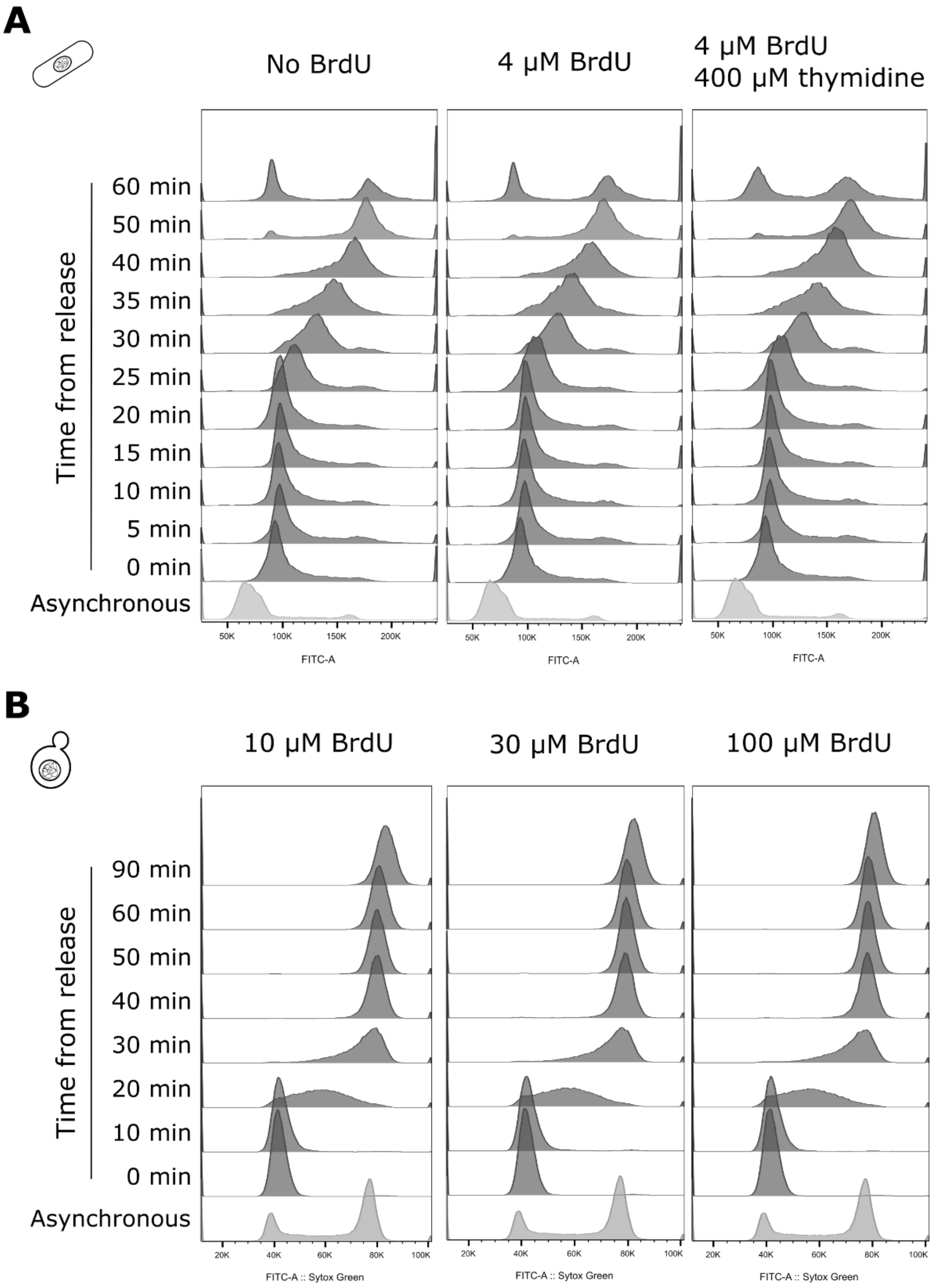


**Supplementary Figure 1: Flow cytometry profiles.** A) Fission yeast flow cytometry profiles. *S. pombe cdc2-as* cells were grown on YES medium and arrested on G2 phase by the addition of 3-BrPP1. Cells were then released synchronously onto the cell cycle by filtering the medium and resuspending in fresh YES medium. BrdU was added 5 minutes after release, and where indicated thymidine was added 30 minutes after release. Before the addition of 3-BrPP1 (asynchronous culture), after the medium exchange (time 0 min), and until the time of culture harvesting (time 60 min), samples were taken for flow cytometry analysis. B) Budding yeast flow cytometry profiles. *S. cerevisiae* cells were grown on YPAD medium and arrested on G1 phase by the addition of alpha factor. Cells were then released synchronously onto one cell cycle by adding pronase two hours after alpha factor addition, and nocodazole 20 min after release. BrdU was added 25 minutes prior to release. Before the addition of alpha factor (asynchronous culture), before the addition of pronase (time 0 min), and until the time of culture harvesting (time 90 min), samples were taken for flow cytometry analysis.

**Supplementary Figure 2**

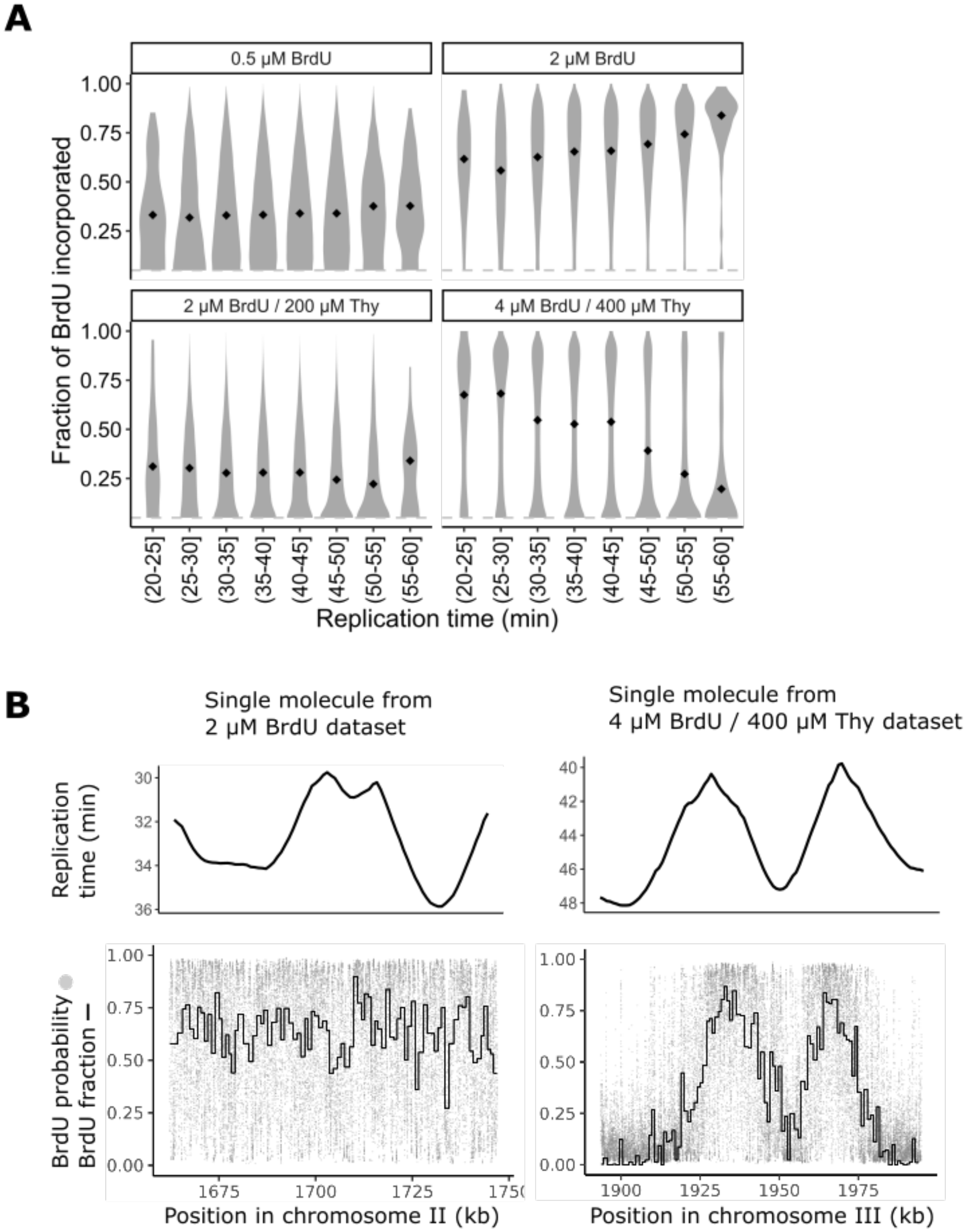


**Supplementary Figure 2: BrdU fraction over replication time from R10 data.** The same DNA samples as shown in Figure 1 were subjected to R10 nanopore sequencing (see Supplementary Table S2). A) BrdU incorporation with respect to replication time in *S. pombe* grown in a range of BrdU and thymidine concentrations. For each single molecule, the fraction of BrdU incorporated in every 1 kb window was determined and plotted in 5 min bins of population level median replication time from (1). The dotted grey line indicates that only BrdU incorporation ≥0.05 are plotted. The black diamonds indicate the median BrdU fraction for each replication time interval. B) Examples of single molecules from the 2 μM BrdU treatment (left) and the 4 μM BrdU / 400 μM thymidine treatment (right). The grey dots indicate the probability of BrdU at each thymidine. The black line indicates the fraction of BrdU in 300 thymidine windows with a probability threshold ≥0.5. The corresponding replication time values are shown on top.

**Supplementary Figure 3**

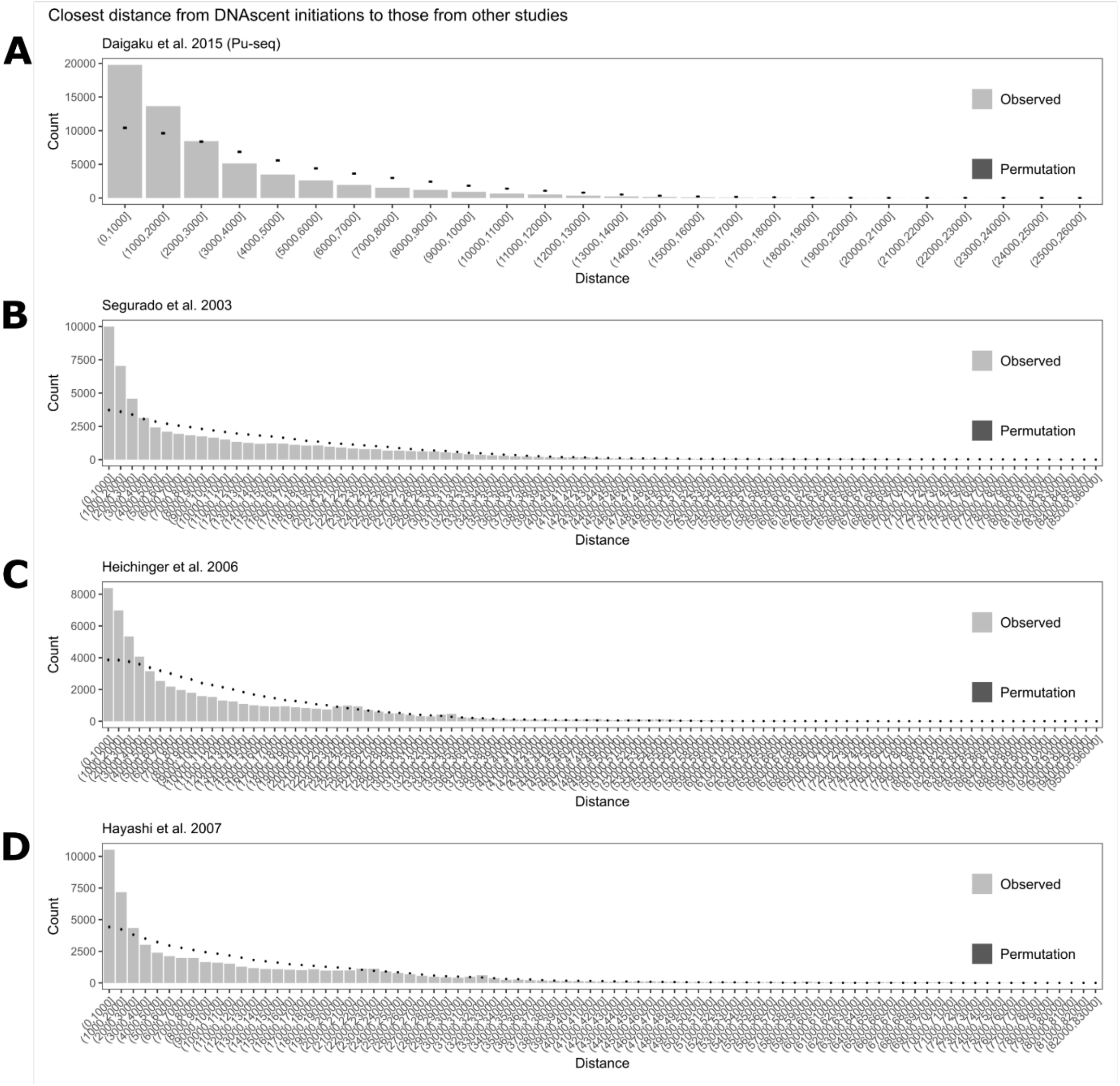


**Supplementary Figure 3. DNAscent initiation sites overlap with initiation sites identified in previous studies.** The distribution of distances between observed DNAscent initiation site midpoints and other datasets are shown as bars. The mean of the counts of distances from a thousand random permutations of the DNAscent initiation genomic intervals are shown as a black dot (standard deviations were computed but are too small to visualise). A) Closest distance from DNAscent initiation sites to Daigaku et al. (Pu-seq) sites (1). B) Closest distance from DNAscent initiation sites to Segurado et al. (2). C) Closest distance from DNAscent initiation sites to Heichinger et al. (3). D) Closest distance from DNAscent initiation sites to Hayashi et al. (4). Note that DNAscent and Pu-seq initiation sites were mapped using the latest reference genome (ASM294v2) which was not available at the time of publication of the other studies.

### Supplementary Figure 4

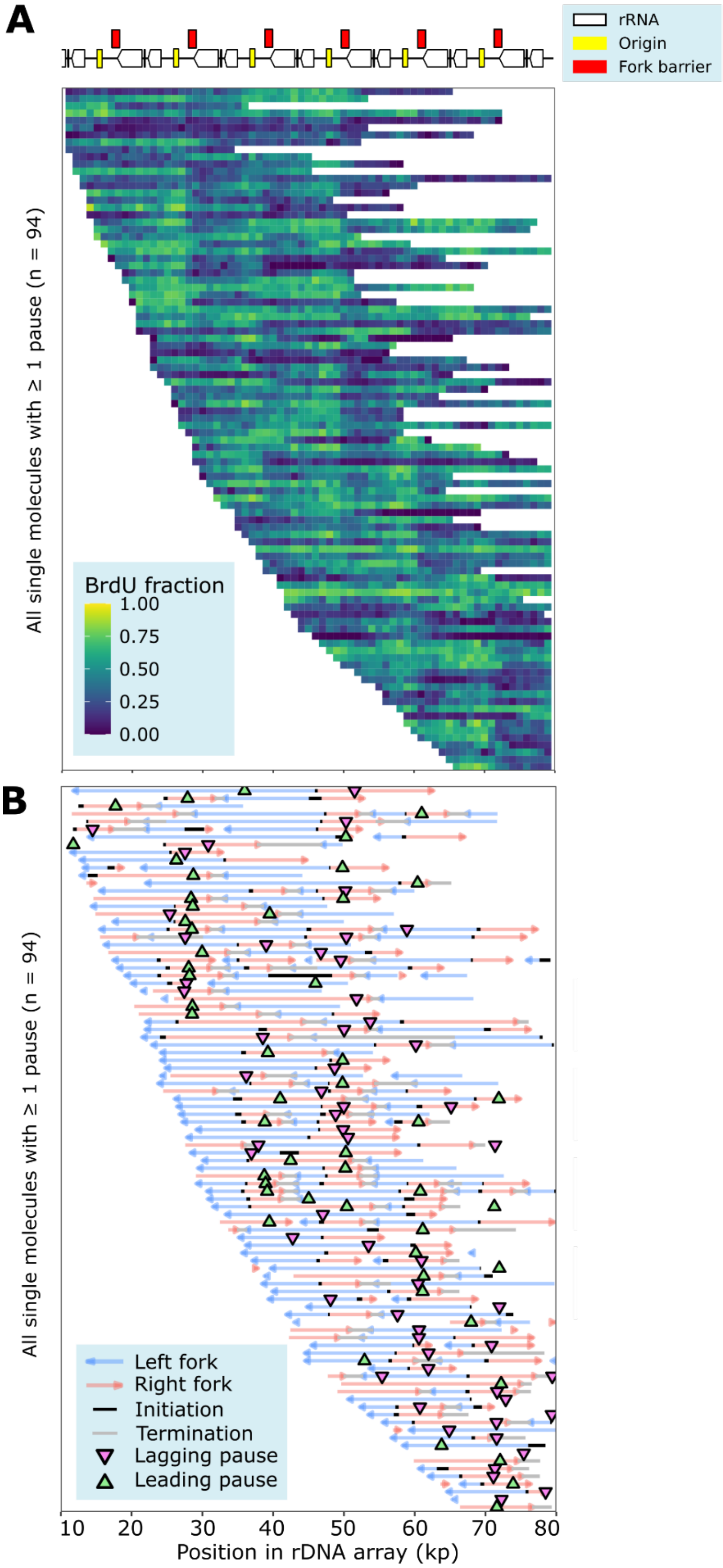

**Supplementary Figure 4. Visualisation of replication dynamics at the rDNA.** A) Heatmap with single molecules with ≥1 replication pause. B) Forks, initiations, terminations, and pauses in the leading or lagging strand in all single molecules from panel B. Note that reads were aligned to a custom rDNA array comprised of 22 copies of rDNA units.

**Supplementary Figure 5**

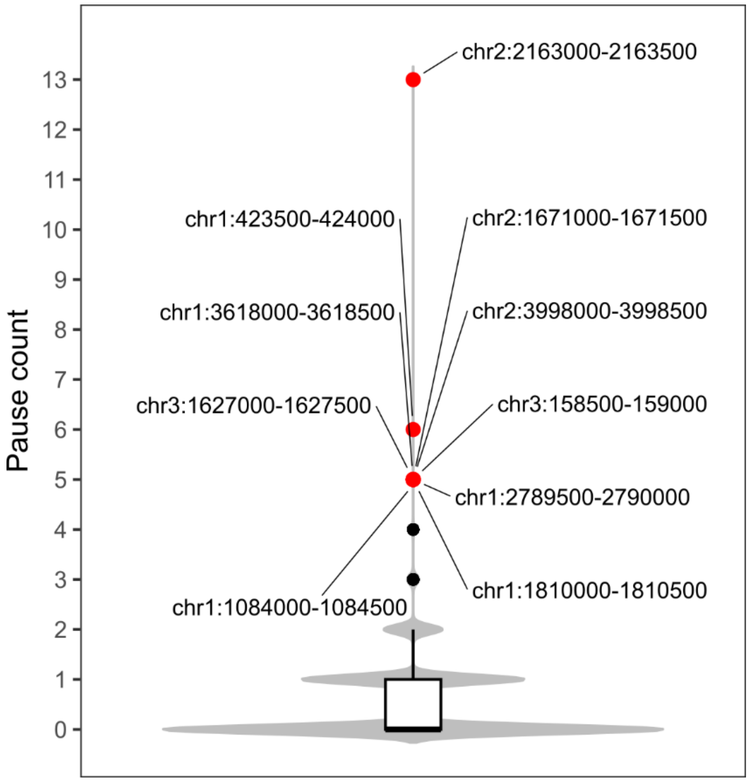


**Supplementary Figure 5. Distribution of pause counts.** Pauses were counted per 500 bp windows using the *de novo* assembly for strain AW2224 as the reference genome. Pause counts ≥5 standard deviations above the mean are highlighted in red and have their coordinates annotated. The highest pause count (coordinates chr2:2163000-2163500) corresponds to the *MPS1* site at the mating-type locus.

**Supplementary Table S1. Biological samples used in this study.**

| Biological_sample | Organism | Strain | BrdU_thymidine_treatment |
|---|---|---|---|
| ARY017_10uM | *Saccharomyces cerevisiae* | ARY017 | 10 µM BrdU |
| ARY017_30uM | *Saccharomyces cerevisiae* | ARY017 | 30 µM BrdU |
| ARY017_100uM | *Saccharomyces cerevisiae* | ARY017 | 100 µM BrdU |
| IDS_43 | *Schizosaccharomyces pombe* | AW2224 | 0.5 µM BrdU |
| IDS_47 | *Schizosaccharomyces pombe* | AW2224 | 2 µM BrdU |
| IDS_49 | *Schizosaccharomyces pombe* | AW2224 | 2 µM BrdU / 20 µM thymidine |
| IDS_52 | *Schizosaccharomyces pombe* | AW2224 | 2 µM BrdU / 200 µM thymidine |
| IDS_63 | *Schizosaccharomyces pombe* | AW2224 | No BrdU |
| IDS_64 | *Schizosaccharomyces pombe* | AW2224 | 4 µM BrdU |
| IDS_65 | *Schizosaccharomyces pombe* | AW2224 | 4 µM BrdU / 400 µM thymidine |

**Supplementary Table S2. Sequencing datasets used in this study.**

| Sequencing_dataset | Biological_sample | Pore_chemistry | Estimated_bases_(Gb) | N50_(kb) | Total_reads_(Million) |
|---|---|---|---|---|---|
| ARY017_10uM-R9 | ARY017_10uM | R9 | 1.74 | 25.06 | 0.168 |
| ARY017_30uM-R9 | ARY017_30uM | R9 | 5.37 | 25.09 | 0.478 |
| ARY017_100uM-R9 | ARY017_100uM | R9 | 7.72 | 23.84 | 0.77 |
| IDS_43-R10 | IDS_43 | R10 | 2.02 | 17.2 | 0.34 |
| IDS_43-R9 | IDS_43 | R9 | 1.94 | 22.86 | 0.178 |
| IDS_47-R10 | IDS_47 | R10 | 2.03 | 21.81 | 0.234 |
| IDS_47-R9 | IDS_47 | R9 | 1.01 | 21.73 | 0.105 |
| IDS_49-R9 | IDS_49 | R9 | 8.8 | 22.94 | 1.06 |
| IDS_52-R10 | IDS_52 | R10 | 2.02 | 22.44 | 0.245 |
| IDS_52-R9 | IDS_52 | R9 | 7.14 | 29.6 | 0.558 |
| IDS_64-R9 | IDS_64 | R9 | 5 | 24.09 | 0.581 |
| IDS_65-R10 | IDS_65 | R10 | 25.81 | 17.37 | 3.28 |
| IDS_65-R9 | IDS_65 | R9 | 17.55 | 20.79 | 2.2 |

**Supplementary Table S3. Files deposited in Zenodo**

| Filename | Sequencing_dataset | Biological_sample | Type_of_file | Alignment_reference |
|---|---|---|---|---|
| ARY017_10uM-R9-SacCer.mod.bam | ARY017_10uM-R9 | ARY017_10uM | mod.bam | SacCer3 |
| ARY017_30uM-R9-SacCer.mod.bam | ARY017_30uM-R9 | ARY017_30uM | mod.bam | SacCer3 |
| ARY017_100uM-R9-SacCer.mod.bam | ARY017_100uM-R9 | ARY017_100uM | mod.bam | SacCer3 |
| IDS_43-R10-ASM294v2.mod.bam | IDS_43-R10 | IDS_43 | mod.bam | ASM294v2 |
| IDS_43-R9-ASM294v2.mod.bam | IDS_43-R9 | IDS_43 | mod.bam | ASM294v2 |
| IDS_47-R10-ASM294v2.mod.bam | IDS_47-R10 | IDS_47 | mod.bam | ASM294v2 |
| IDS_47-R9-ASM294v2.mod.bam | IDS_47-R9 | IDS_47 | mod.bam | ASM294v2 |
| IDS_49-R9-ASM294v2.mod.bam | IDS_49-R9 | IDS_49 | mod.bam | ASM294v2 |
| IDS_52-R10-ASM294v2.mod.bam | IDS_52-R10 | IDS_52 | mod.bam | ASM294v2 |
| IDS_52-R9-ASM294v2.mod.bam | IDS_52-R9 | IDS_52 | mod.bam | ASM294v2 |
| IDS_64-R9-ASM294v2.mod.bam | IDS_64-R9 | IDS_64 | mod.bam | ASM294v2 |
| IDS_65-R10-ASM294v2.mod.bam | IDS_65-R10 | IDS_65 | mod.bam | ASM294v2 |
| IDS_65-R9-ASM294v2.mod.bam | IDS_65-R9 | IDS_65 | mod.bam | ASM294v2 |
| IDS_65-R9-AW2224_assembly.mod.bam | IDS_65-R9 | IDS_65 | mod.bam | AW2224_assembly |
| IDS_65-R9-rDNA.mod.bam | IDS_65-R9 | IDS_65 | mod.bam | rDNA |
| reference_AW2224_assembly.fasta | IDS_65-R9 | IDS_65 | fasta | Not_applicable |
| reference_rDNA.fasta | IDS_65-R9 | IDS_65 | fasta | Not_applicable |
| DNAscent_initiation_sites-IDS_65-R9-ASM294v2.bedgraph | IDS_65-R9 | IDS_65 | bedgraph | ASM294v2 |
| DNAscent_termination_sites-IDS_65-R9-ASM294v2.bedgraph | IDS_65-R9 | IDS_65 | bedgraph | ASM294v2 |
| DNAscent_leftward_forks-IDS_65-R9-ASM294v2.bedgraph | IDS_65-R9 | IDS_65 | bedgraph | ASM294v2 |
| DNAscent_initiation_sites-IDS_65-R9-AW2224_assembly.bedgraph | IDS_65-R9 | IDS_65 | bedgraph | AW2224_assembly |
| DNAscent_termination_sites-IDS_65-R9-AW2224_assembly.bedgraph | IDS_65-R9 | IDS_65 | bedgraph | AW2224_assembly |
| DNAscent_leftward_forks-IDS_65-R9-AW2224_assembly.bedgraph | IDS_65-R9 | IDS_65 | bedgraph | AW2224_assembly |

| | | | | |
|---|---|---|---|---|
| DNAscent_pause_sites-IDS_65-R9-AW2224_assembly.bedgraph | IDS_65-R9 | IDS_65 | bedgraph | AW2224_assembly |
| DNAscent_initiation_sites-IDS_65-R9-rDNA.bedgraph | IDS_65-R9 | IDS_65 | bedgraph | rDNA |
| DNAscent_termination_sites-IDS_65-R9-rDNA.bedgraph | IDS_65-R9 | IDS_65 | bedgraph | rDNA |
| DNAscent_leftward_forks-IDS_65-R9-rDNA.bedgraph | IDS_65-R9 | IDS_65 | bedgraph | rDNA |
| DNAscent_leading_pause_sites-IDS_65-R9-rDNA.bedgraph | IDS_65-R9 | IDS_65 | bedgraph | rDNA |
| DNAscent_lagging_pause_sites-IDS_65-R9-rDNA.bedgraph | IDS_65-R9 | IDS_65 | bedgraph | rDNA |
| replication_time_s_cerevisiae.bed | Not_applicable | Not_applicable | bed | SacCer |
| replication_time_s_pombe.bed | Not_applicable | Not_applicable | bed | ASM294v2 |
| annotation_centromeres_AW2224_assembly.bed | Not_applicable | Not_applicable | bed | AW2224_assembly |
| annotation_mating-type_locus_AW2224_assembly.bed | Not_applicable | Not_applicable | bed | AW2224_assembly |
